# Characterization of SARS-CoV-2 ORF6 deletion variants detected in a nosocomial cluster during routine genomic surveillance, Lyon, France

**DOI:** 10.1101/2020.08.07.241653

**Authors:** Grégory Quéromès, Grégory Destras, Antonin Bal, Hadrien Regue, Gwendolyne Burfin, Solenne Brun, Rémi Fanget, Florence Morfin, Martine Valette, Bruno Lina, Emilie Frobert, Laurence Josset

## Abstract

Through routine genomic surveillance of the novel SARS-CoV-2 virus (n=229 whole genome sequences), 2 different frameshifting deletions were newly detected in the open reading frame (ORF) 6, starting at the same position (27267). While the 26-nucleotide deletion variant was only found in one sample in March 2020, the 34-nucleotide deletion variant was found within a single geriatric hospital unit in 5/9 patients sequenced and one health care worker with samples collected between April 2^nd^ and 9^th^, 2020. Both the presence of the 34-nucleotide deletion variant limited to this unit and the clustering of the corresponding whole genome sequences by phylogeny analysis strongly suggested a nosocomial transmission between patients. Interestingly, prolonged viral excretion of the 34-nucleotide deletion variant was identified in a stool sample 14 days after initial diagnosis for one patient. Clinical data revealed no significant difference in disease severity between patients harboring the wild-type or the 34-nucleotide deletion variants. The *in vitro* infection of the two deletion variants on primate endothelial kidney cells (BGM) and human lung adenocarcinoma cells (Calu-3) yielded comparable replication kinetics with the wild-type strain. Furthermore, high viral loads were found *in vivo* regardless of the presence or absence of the ORF6 deletion. Our study highlights the transmission and replication capacity of two newly described deletion variants in the same ORF6 region.

**Importance:** While the SARS-CoV-2 genome has remained relatively stable since its emergence in the human population, genomic deletions are an evolutionary pattern previously described for the related SARS-CoV. Real-time genomic monitoring of the circulating variants is paramount to detect strain prevalence and transmission dynamics. Given the role of ORF6 in interferon modulation, further characterization, such as mechanistic interactions and interferon monitoring in patients, is crucial in understanding the viral-host factors driving disease evolution.

## Introduction

The coronavirus disease 2019 (COVID-19) pandemic triggered by the novel severe acute respiratory syndrome coronavirus 2 (SARS-CoV-2) virus has continued to spread globally since its emergence in China in late 2019 (1, 2). Countries and localities have implemented various levels of public health mitigation measures with debatable success in an effort to control virus propagation (3–6). The challenge in better understanding the fundamental characteristics of this novel virus includes the heterogeneous disease reports in conjunction with no clear treatments or vaccines yet available or approved (7–9). Epidemiological tracking is paramount in the context of this current pandemic (10, 11). In particular, the genomic surveillance of circulating virus variants, such as with the seasonal epidemics of the influenza virus or even with the 2003 SARS epidemic, has brought useful information in understanding their respective evolutionary dynamics (12, 13). Recent tracking reports have discussed the high frequency and global distribution of a variant harboring the D614G substitution located on the SARS-CoV-2 spike protein. While higher infectious titer and increased protein stability have been associated with this variant, a clear fitness advantage has not been unequivocally established (14, 15). Historically, evolution of the related SARS-CoV virus is defined by deletion regions that impact the open-reading frames (ORF) of its genome (16, 17). Several deletions of large variations in size and prevalence have already been described in the SARS-CoV-2 genome (18–20).

The aim of this study was to therefore describe clinical patient data and the viral replication capacity of two newly detected ORF6 deletion variants detected in early April from routine genomic surveillance of COVID-19 patients in Lyon, France.

## Results

### ORF6 Deletion Variants Detected During Routine Genomic Surveillance

As part of the Auvergne-Rhône-Alpes (ARA) regional surveillance, 229 samples collected between Feb 2^nd^ and April 12^th^ were sequenced by the French National Reference Center of Respiratory Viruses. These samples originated mainly from the Hospices Civils de Lyon (HCL) (149 sequences from 58 units within 11 different hospital sites), with some from other hospitals in the Lyon area (24 sequences) and other regional hospitals (56 sequences, 12 cities).

Of these 229 samples, 6 sequences were shown to carry a 34-nt deletion (at position 27267-27300), henceforth denoted as D34, and 1 sequence a 26-nt deletion (at position 27267-27292), denoted as D26 (Figure 1). These deletions are both frameshifting deletions in the ORF6, starting at the same 27267 position after a stretch of 3 T at 27264-27266 (Figure 2). Because of the frameshifting, the D34 variant generates a premature stop codon (at position 27308, Wuhan-Hu-1 numbering), resulting in a presumed truncated 24 amino acid protein, instead of 61 in the wild-type (Wuhan-Hu-1 and all other sequences described yet). The D26 variant yields a 28 amino acid protein with its premature stop codon at position 27312 (Wuhan-Hu-1 numbering) (Figure 2). These deletions have not yet been described on the CoV-GLUE resource, which lists all genomic variants in SARS-CoV-2 sequences available on the GISAID database (Supplementary Data 1). Of note, ORF6 variants are annotated at different positions in CoV-GLUE to maximize amino acid alignment.

**FIGURE 1.**
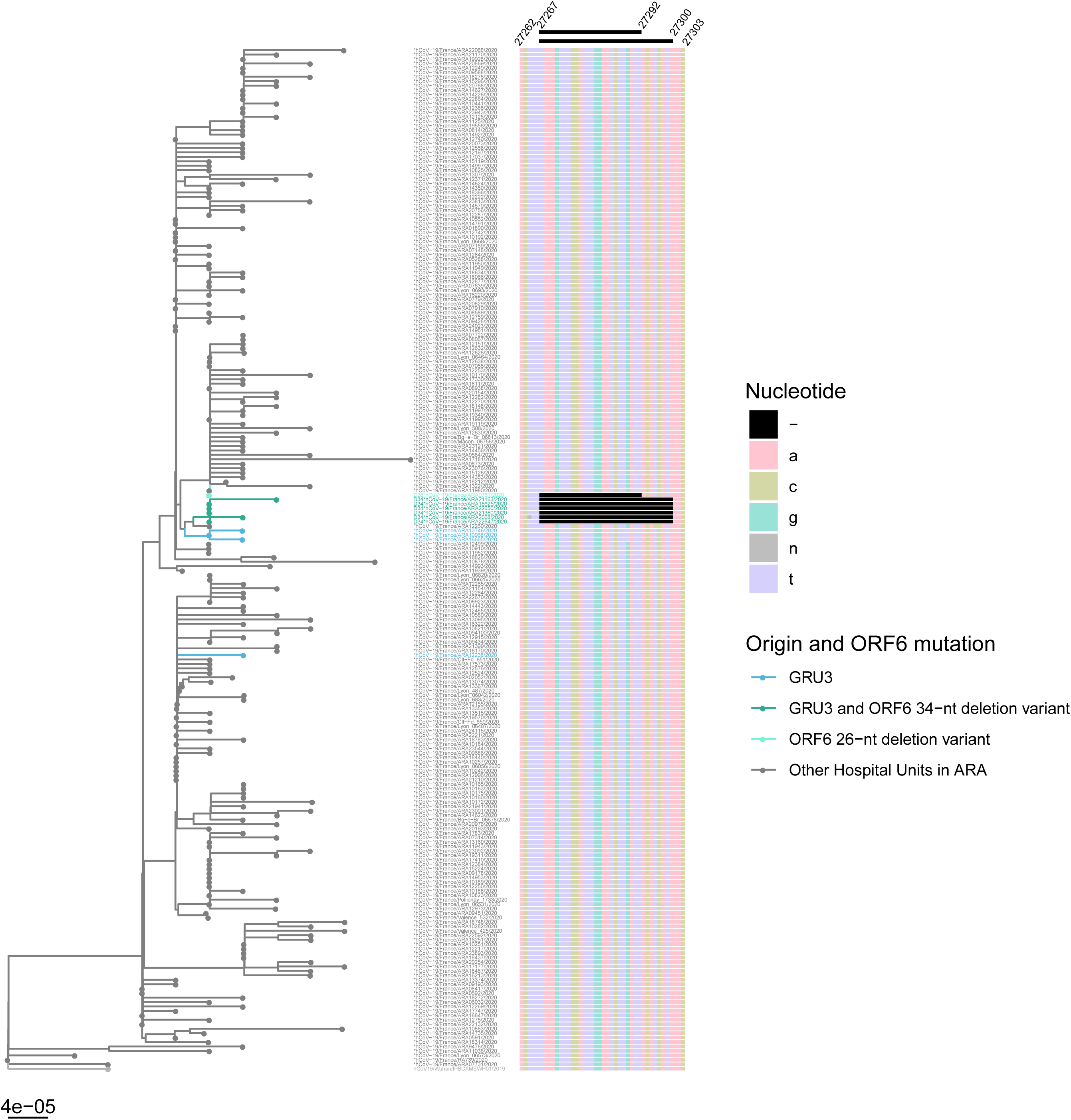
Phylogenetic tree of SARS-CoV-2 full genome sequences from ARA patients (n=229). The phylogenetic tree was constructed using R software using ape package and the neighbor joining evolutionary method (hCoV19/Wuhan/IPBCAMSWH01/2019 as the root). The colored branches denote the hospital unit origin of the sequence and ORF6 status. On the right, multiple sequence alignment from nucleotide position 27267-27303 (Wuhan-Hu-1 numbering) is illustrated, with the 26-nt and 34-nt deletions depicted in black. The deletion sites of interest were not included for genetic distance calculation.

**FIGURE 2.**
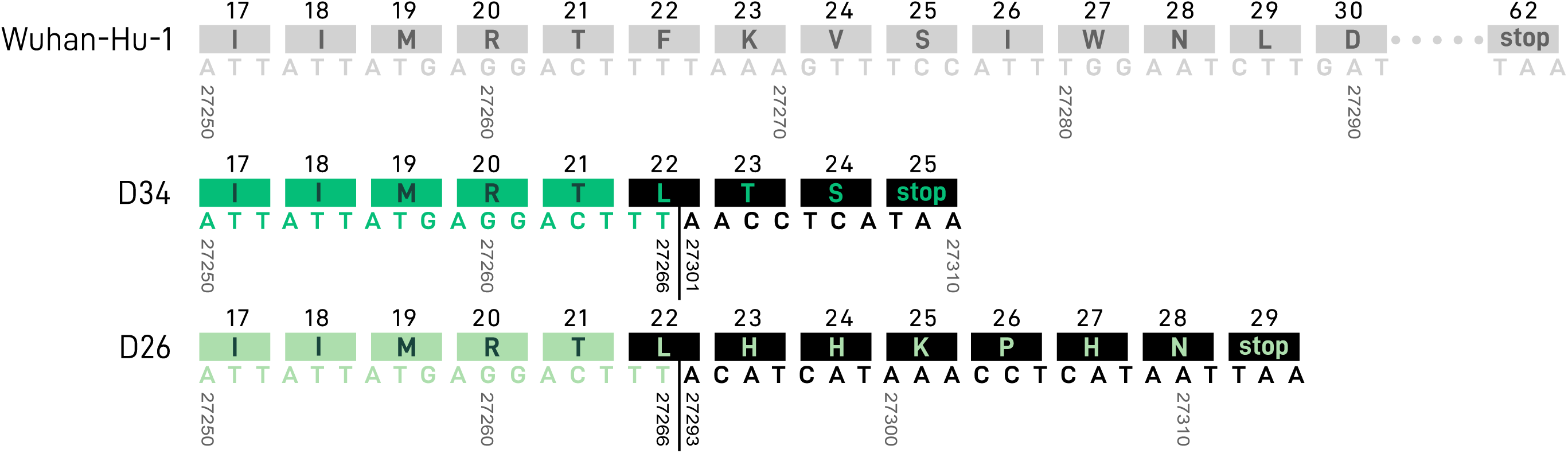
SARS-CoV-2 genome map at ORF6 position 27250-27290. Both 34-nt (D34) and 26-nt (D26) deletion regions are illustrated in green, with the Wuhan-Hu-1 reference genome in grey. Amino acids are depicted in the colored blocks above the corresponding nucleic acids. Nucleic acid and amino acid alterations resulting from the ORF6 deletions are illustrated in a black. Image adapted from the genome visualization tool from the CoV-GLUE online resource, enabled by data from GISAID.

The 7 sequences carrying an ORF6 deletion belong to lineage B1, a lineage widely circulating in Europe (Supplementary Data 1). There were between 0 and 3 SNP (Single Nucleotide Polymorphism) differences among D34 strains, for which 3/6 mutants displayed 1 to 3 SNPs, and between 2 and 4 SNPs between D26 and D34 strains (Supplementary Data 2).

### Evidence for direct transmission of the ORF6 34-nt deletion variant

The D34 samples were all collected from hospitalized patients or health care workers (HCW) in the same geriatric rehabilitation unit in the Hospices Civils de Lyon (GRU-3), between April 2nd and April 9^th^, while the sample with the 26-nt deletion was collected one month earlier (March 10^th^) in a geriatric unit of another hospital (Table 1). The hospitals are 80 km apart and there was no evidence for the transfer of patient #73 with the 26-nt deletion into GRU-3. To track the origin of the deletion, all patients hospitalized in the GRU-3 geriatric unit and all samples collected between March 18^th^ and April 9^th^ with high viral loads of SARS-CoV-2 (RT-PCR Ct value <20) were sequenced (n=9). In total, 44% (4/9 patients) were infected with the WT SARS-CoV-2 ORF6 (samples collected between March 18^th^ −30^th^), with no read carrying the deletion at a minor frequency. Out of the 4 WT SARS-CoV-2 ORF6, three sequences were very similar to D34 and carried a G27289T SNP (D30Y), which has already been identified in three patients from England between April 20^th^ and 27^th^, 2020 (Figure 1). The other 55% (5/9 patients), in addition to 1 HCW, were infected with the 34-nt deletion (samples collected after April 2^nd^) with 100% of the reads carrying the deletion for each patient. Overall, 8/9 sequences of GRU-3, corresponding to those of the D34 variants and those of the three WT strains carrying the ORF6 G27289T SNP were clustered together, while the sequence of the patient #38 was more divergent. We could not investigate whether the mutation spread after April 9^th^ as only one COVID-19 patient was hospitalized in this geriatric unit since, for which their viral load (Ct>30) was too low for mNGS.

**TABLE 1.** Clinical data from patients infected with the ORF6 deletion variants compared with related patients infected with WT SARS-CoV-2 strains

|  | Patient #68 | Patient #38 | Patient #46 | Patient #65 | Patient #25 | Patient #8 | HCW#63 | Patient #60 | Patient #47 | Patient #50 |
| --- | --- | --- | --- | --- | --- | --- | --- | --- | --- | --- |
| **Sequence name<sup>a</sup>** | ARA10968/2020 | ARA12238/2020 | ARA13746/2020 | ARA16665/2020 | ARA18625/2020 | ARA2608/2020 | ARA21163/2020 | ARA21360/2020 | ARA22647/2020 | ARA22650/2020 |
| **Age (years)** | 89 | 78 | 84 | 89 | 97 | 82 | 31 | 94 | 81 | 85 |
| **Sex** | F | M | F | F | M | F | F | F | M | F |
| **Comorbidities** | Hypertension | None | None | Cardiovascular disease, Hypertension | Cardiovascular disease, Hypertension | Cardiovascular disease, Hypertension, Obesity | NA | Cardiovascular disease, Hypertension | None | Cardiovascular disease |
| **Respiratory manifestations** | URTI | LRTI | LRTI | Asymptomatic | Asymptomatic | LRTI | URTI | URTI | Not specified | LRTI |
| **Hospitalization (Yes / No)** | Yes | Yes | Yes | Yes | Yes | Yes | No | Yes | Yes | Yes |
| **Hospitalization Unit** | GRU-3 | GRU-3 | GRU-3 | GRU-3 | GRU-3 | GRU-3 | GRU-3 | GRU-3 | GRU-3 | GRU-3 |
| **ORF6 deletion** | None | None | None | None | D34 | D34 | D34 | D34 | D34 | D34 |
| **Date of diagnostic** | 2020/03/18 | 2020/03/22 | 2020/03/25 | 2020/03/30 | 2020/04/02 | 2020/04/06 | 2020/04/07 | 2020/04/07 | 2020/04/09 | 2020/04/09 |
| **Duration of excretion<sup>b</sup>** | Respiratory sample positive at D21 | Not monitored | Not monitored | Respiratory sample positive at D14 | Not monitored | Not monitored | Not monitored | Not monitored | Respiratory and stool samples positive at D14 | Not monitored |
| **Outcome<sup>b</sup>** | Favorable | Favorable | Favorable | Favorable | Deceased, unrelated to COVID-19 (D5) | Deceased from COVID-19 (D6) | Favorable | Favorable | Favorable | Deceased from COVID-19 (D9) |

HCW=health care worker; COPD=chronic obstructive pulmonary disease; URTI: Upper Respiratory Tract Infection; LRTI: Lower Respiratory Tract Infection; HCL=Hospices Civils de Lyon; GRU-1=geriatric rehabilitation unit within HCL; D26=26-nt deletion; D34=34-nt deletion; D=Day
<sup>a</sup> Sequence names in the table are preceded by hCoV-19/France/; example: hCoV-19/France/ARA0731/2020
<sup>b</sup> Duration of excretion and death are calculated from the date of diagnosis

### 34-nt ORF6 deletion variants yielded similar clinical presentations as WT ORF6 in hospitalized patients

Clinical data were studied on hospitalized patients in GRU-3 (i.e. excluding the HCW #63 to better control for confounding variables (e.g. age and comorbidities)) to compare COVID-19 severity between patients infected with D34 and with WT SARS-CoV-2 (n=9) (Table 1). The median age of hospitalized patients was 87 years (ranging from 78 - 97), with 7 patients presenting at least cardiovascular disease as a risk factor (Table 1). Other comorbidities included hypertension (n=5), obesity (n=1), and chronic obstructive pulmonary disease (n=1).

Clinical presentations of hospitalized patients with the D34 variants (n=5) were classified as asymptomatic for one patient, upper respiratory tract infection for 2 patients, and lower respiratory tract infection (pneumonia) for 3 patients. To evaluate disease severity in relation to the D34 deletion, mild (asymptomatic and URTI, n=5) versus severe COVID-19 (LRTI, n=4) was compared by Fisher’s exact test. No significant difference in clinical presentation could be observed between hospitalized patients harboring or not the ORF6 deletion (p>0.99).

From the five hospitalized patients harboring D34 deletion, 2 died from COVID-19 infection, all presenting LRTI and comorbidities. One patient (#25) died at day 5 after diagnosis, but their death was not related to COVID-19 infection but to septicemia. To evaluate disease outcome in relation to the D34 deletion, death from COVID-19 versus favorable outcome (including non-COVID-19 death) was compared by Fisher’s exact test. No significant difference in disease outcome could be observed between hospitalized patients harboring or not the D34 deletion (p=0.44).

Notably, patient #47 harboring a D34 variant was still positive after 14 days in respiratory and stool samples. Virus present in the stool was 100% identical to the first virus sequenced from respiratory samples. Unfortunately, the respiratory sample at day 14 could not be sequenced due to Ct >30.

### SARS-CoV-2 deletion variants yield comparable replication kinetics to reference strain

Two genomes representative of ORF6 deletion variants found in this regional circulation were selected for replication tests: hCoV-19/France/ARA22647/2020 (D34 variant) and hCoV-19/France/ARA0731/2020 (D26 variant). These genomes were compared against the reference genome hCoV-19/France/ARA24023/2020 (devoid of any deletions), which was the most similar isolated strain sequenced available in the laboratory at the time of the investigation. The reference genome had 1 to 3 SNPs compared with D26 and D34 variants (Supplementary Data 2).

Replication kinetics measured by viral genome quantification revealed no significant difference between the three strains throughout the course of *in vitro* infection on both BGM and Calu-3 cell lines (Figure 3). However, a significant difference was observed between cell lines for each strain, with an increased level of replication on BGM (as early as 24 hours post-inoculation). More specifically, viral replication spiked rapidly on BGM cells within the first 48 hours, before reaching a plateau at 72 hours. Conversely, viral replication on Calu-3 cells rose steadily within the first 48 hours, before reaching a plateau at 96 hours. Of interest, a 2-log difference was observed for maximum genome quantification between BGM and Calu-3, with an average of 5.76×10^12^ and 4.01×10^10^ copies/mL, respectively.

**FIGURE 3.**
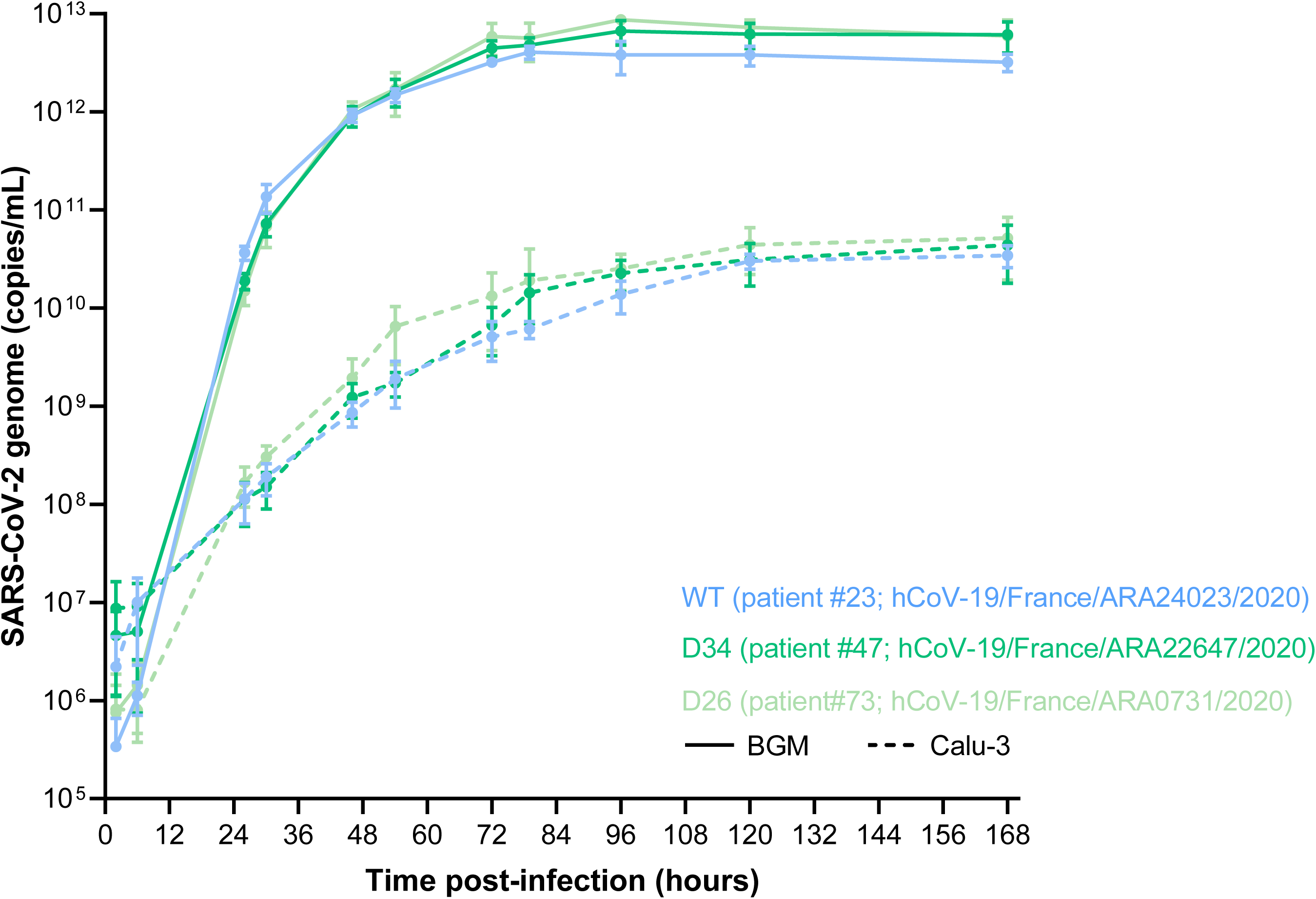
Replication kinetics of SARS-CoV-2 ORF6 deletion variants on BGM and Calu-3 cell lines. SARS-CoV-2 strains were inoculated on confluent BGM and Calu-3 cells at an MOI of 10^−3^ and then incubated at 36°C with 5% CO_2_ for 7 days. Supernatant samples were collected at regular intervals, for which viral particle quantification was performed by qRT-PCR. Each data point is the average of three replicates, with standard deviation as error bars. Statistical analysis was performed by two-way ANOVA with Tukey multiple comparisons between both factors of comparison (virus strain and cell line) on GraphPad Prism (software version 8.4.3).

## Discussion

Despite reports of the relative stability of the SARS-CoV-2 genome within the human population, whole genome sequencing has revealed recurrent variants with variable mutation patterns over the course of the pandemic and within distinct geographic regions (10, 18, 21, 22). Here we describe clinical data and the viral replication ability associated with large ORF6 deletion variants identified through surveillance of patients from the same hospital unit. The CoV-GLUE phylogenetic resource revealed no other international sequence harboring the ORF6 deletions described herein. However, other reports of similar patterns of genomic deletions in the SARS-CoV-2 genome since its emergence have already been described, including in the ORFs 6, 7 and 8 (19, 20, 22, 23).

The origin of the D34 deletion is still unknown. However, as the WT virus isolated from GRU-3 patients were genetically close to the D34 variants, the GRU-3 patients infected by the WT virus in March could have been the initial source of the D34 deletion. Nevertheless, its introduction since April 2^nd^ with its limited presence in the hospital unit thereafter and the clustering of the corresponding whole genome sequences by phylogeny analysis strongly suggest a nosocomial transmission of the D34 variant. Importantly, the persistence of the same D34 consensus sequence in a patient’s stool sample 14 days after diagnosis from a nasopharyngeal sample gives emphasis to the enteric tropism capacity of D34 variants and the potential contribution to nosocomial transmission (24). None of the GRU-3 patients with WT or deletion strains presented any intra-host diversity in the ORF6 deletion region that would have been indicative of a recent mutation or recombination.

The similar consensus sequences in the nasopharyngeal and in the stool samples over a 2-week period suggest an apparent low intra-host evolution. However, different sites of SARS-CoV-2 replication might have a different impact on the intra-host evolution, but we could not provide the consensus sequence from the nasopharyngeal sample collected at day 14 due to low viral load to confirm this hypothesis (25). Moreover, the normal rate of mutation of SARS-CoV-2 has been reported at about 2.5 mutations per month (26, 27). Nevertheless, the fact that D34 variants had 1 to 3 SNP differences between consensus sequences of D34 strains collected between one week might highlight a higher mutation rate than normal and be linked to adaptative mutations following the deletion. Evidence of adaptation by means of genomic deletions during the middle and late phases of the SARS-CoV 2003 epidemic has been tenuously described (16, 17, 28–30). Our restrictive number of sequenced strains to date does not currently allow conclusive population prevalence information for the Auvergne-Rhône-Alpes region. The importance of such genomic variants by NGS investigation during the evolution of disease transmission and population prevalence should not be overlooked (13, 31, 32). Further genomic surveillance is needed to assess specific evolutionary patterns of these variants.

Our study characterized the D34 and D26 ORF6 variants, showing no significant impact on replication *in vitro* in comparison to a wild-type strain, in two different cell lineages. The comparable replication kinetics between wild-type and deletion strains determined *in vitro* and *in vivo* replication capacity (the latter being assessed by RT-PCR from diagnosis, Ct<20) is supported by the congruent in vivo replication capacity (the latter being assessed by RT-PCR from diagnosis, Ct<20). Furthermore, there was no significant difference in disease severity between patients at the GRU-3 hospital unit harboring D34 ORF6 variant or WT. It is important to note that interferon immunoprofiling of patients, a key factor in disease manifestation, was not assessed and remains to be explored (33).

Research on the SARS-CoV ORF6 has attributed this accessory protein with potential functions of intracellular membrane rearrangements, of interferon induction inhibition, and of replication stimulation (34–36). Recent literature confirms the interferon signaling inhibitory function of the SARS-CoV-2 ORF6 protein (37). Another recent study reported a 27-nt in-frame ORF6 deletion (at position 27264 - 27290) and demonstrated important three-dimensional structural alterations to the protein (23). Whereas this in-frame deletion variant would have emerged during passaging on VeroE6 cells, the D34 and D26 variants already presented these deletions in the initial clinical isolates before cell culture. Frameshift modifications, especially the −1 programmed ribosomal frameshift allowing ORF1b translation, have been known to alter coronavirus genomic and subgenomic RNA production efficiencies (38–41). And so, we can postulate that ORF6-mediated frameshifts can affect downstream elements, such as the critically multifunctional nucleocapsid (N) protein (42, 43). Further genomic and structural investigations are needed to explore the impact of these ORF6 deletions, in terms of ribosomal frameshift stimulators and RNA translation production ratios, as well as innate host immunity modulation. The integration of more fundamental research dedicated to elucidating the factors that impact SARS-CoV-2 replication, transmission, and disease progression will ultimately help translational projects to advance the fight against the current COVID-19 pandemic.

## Material and Methods

### Sequencing

Early routine genomic surveillance of SARS-CoV-2 in the National Reference Center (NRC) of Respiratory Viruses is based on daily random selection of samples with SARS-CoV-2 detected with quantitative reverse-transcriptase polymerase chain reaction (qRT-PCR) cycle threshold (Ct) <20 (6), which were then sequenced using an RNA metagenomic next-generation sequencing (mNGS) method previously described (18). Briefly, viral genetic material contained in nasopharyngeal and stool samples was extracted by the EMAG® platform (bioMerieux, Lyon, FR). After DNAse treatment (Life Technologies, Carlsbad, CA, USA), samples underwent random amplification using Whole Transcriptome Amplification (WTA2 kit, Sigma-Aldrich, Darmstadt, DE) before sequencing on an Illumina NextSeqTM 550 with mid-output 2×150 flow cell. Importantly, the strains displaying an ORF6 deletion were confirmed by 3 other techniques, including capture- and amplicon-based strategies (44). Sequencing of patient samples began on Feb 8^th^ and is ongoing. For the stool sample, an amplicon-based approach developed by the ARTIC network (https://artic.network/ncov-2019) combined with Oxford Nanopore Technologies sequencing was used.

### Phylogeny

Multiple sequence alignment was performed using the DECIPHER package in R (45). Pairwise distances were computed using the Kimura (K80) model implemented in the function dist.dna, deleting the sites with missing data in a pairwise way. The phylogenetic tree was constructed using R software using ape package and the neighbor joining evolutionary method (hCoV19/Wuhan/IPBCAMSWH01/2019 as the root). CoV-GLUE resource [http://cov-glue.cvr.gla.ac.uk, (46)] was used to generate phylogenetic placement of the mutants, annotate the sequences, and check the prevalence of the deletions among worldwide sequences. Codon numbering is based on the Wuhan-Hu-1 sequence.

### Virus replication kinetics

Replication kinetics was performed on both confluent buffalo green monkey (BGM) (BioWhittaker Europe) and human lung adenocarcinoma (Calu-3) cells (ATCC® HTB-55™, Plateforme iPS, NeuroMyoGene Institute, Lyon, FR) at a multiplicity of infection (MOI) of 10^−3^ at 37°C for 7 days, fully respecting the WHO interim biosafety guidance related to the coronavirus disease (47). Comparative viral particle quantification of culture supernatant was performed by RdRp Institut Pasteur qRT-PCR on a QuantStudio™ 5 System (Applied Biosystems, ThermoFischer Scientific) with a standard curve after semi-automated EMAG® extraction (bioMérieux, Lyon, FR) (6). Statistical analysis was performed by two-way ANOVA with Tukey multiple comparisons between both factors of comparison (virus strain and cell line) on GraphPad Prism (software version 8.4.3).

### Ethics

Samples used in this study were collected as part of an approved ongoing surveillance conducted by the National Reference Center for Respiratory Viruses (NRC) in France (WHO reference laboratory providing confirmatory testing for COVID-19). The investigations were carried out in accordance with the General Data Protection Regulation (Regulation (EU) 2016/679 and Directive 95/46/EC) and the French data protection law (Law 78–17 on 06/01/1978 and Décret 2019–536 on 29/05/2019). Samples were collected for regular clinical management during hospital stay, with no additional samples for the purpose of this study. Patients were informed of the research and their non-objection approval was confirmed. This study was presented by the ethics committee of the Hospices Civils de Lyon (HCL), Lyon, France and registered on the HCL database of RIPHN studies (AGORA N°41).

## Data availability

The SARS-CoV-2 genomes sequenced in this study were deposited on the GISAID database (https://www.gisaid.org/) on a regular basis, accession numbers can be found in Supplementary Table 1.

## Acknowledgments

We would like to thank all the patients, laboratory technicians, and clinicians who contributed to this investigation. We gratefully acknowledge all the members of CoV-GLUE, Nextstrain.org, and virological.org for sharing their analysis in real time.

## Competing Interests

The authors declare no competing interests.

BL is a member of the French Scientific Committee for SARS-CoV-2.

---

**SUPPLEMENTARY FIGURE 1.**
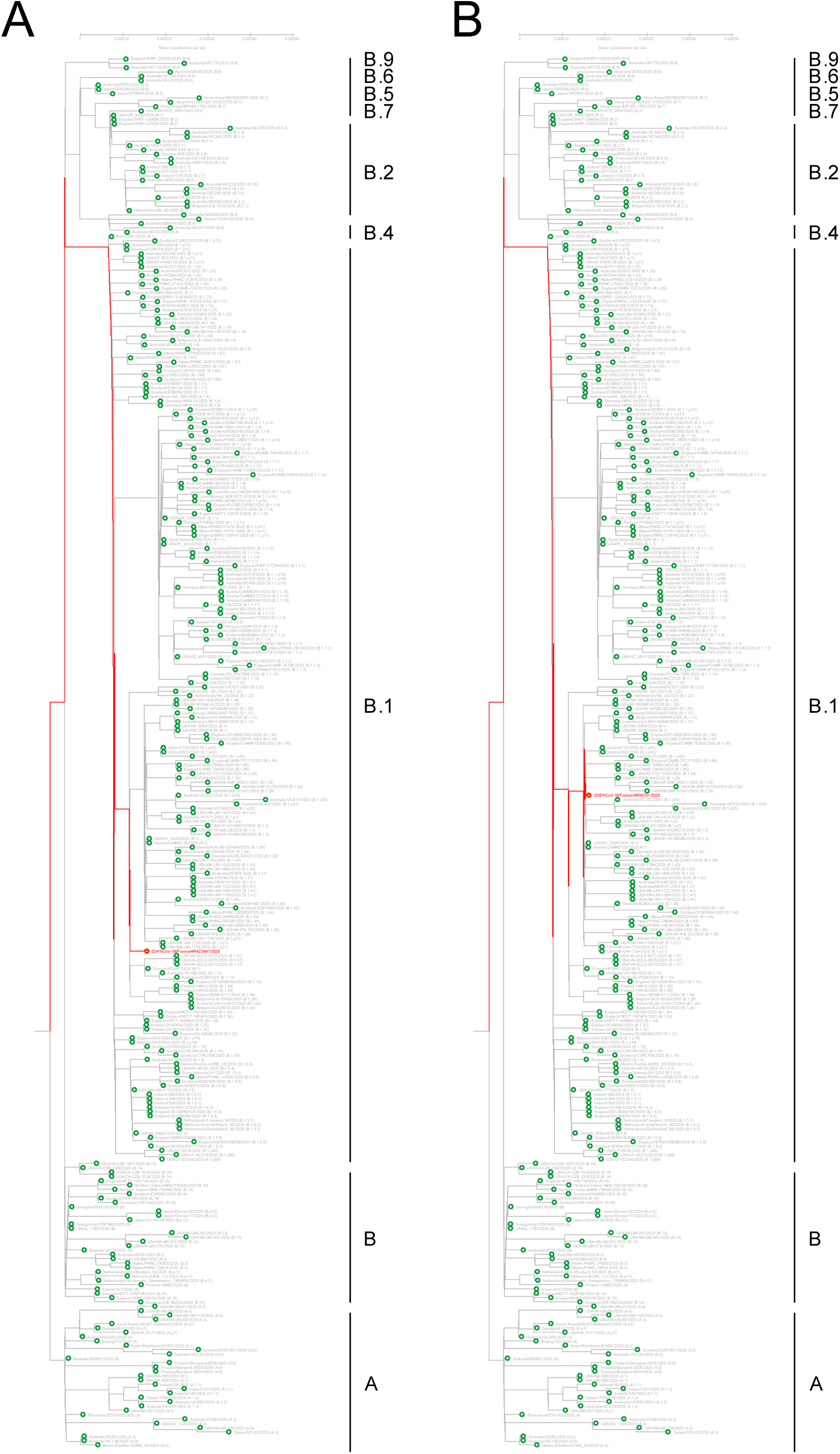
CoV-GLUE phylogenomic placement map. The D34 (A) and D26 (B) deletions are phylogenetically positioned against global SARS-CoV-2 sequences deposited on the GISAID database and annotated with PANGOLIN lineages. The tree was generated by the CoV-GLUE resource, which uses the RAxML (Randomized Axelerated Maximum Likelihood) software (Stamatakis 2014). Relevant deletions are in red, while WT sequences are in green.

**SUPPLEMENTARY FIGURE 2.**
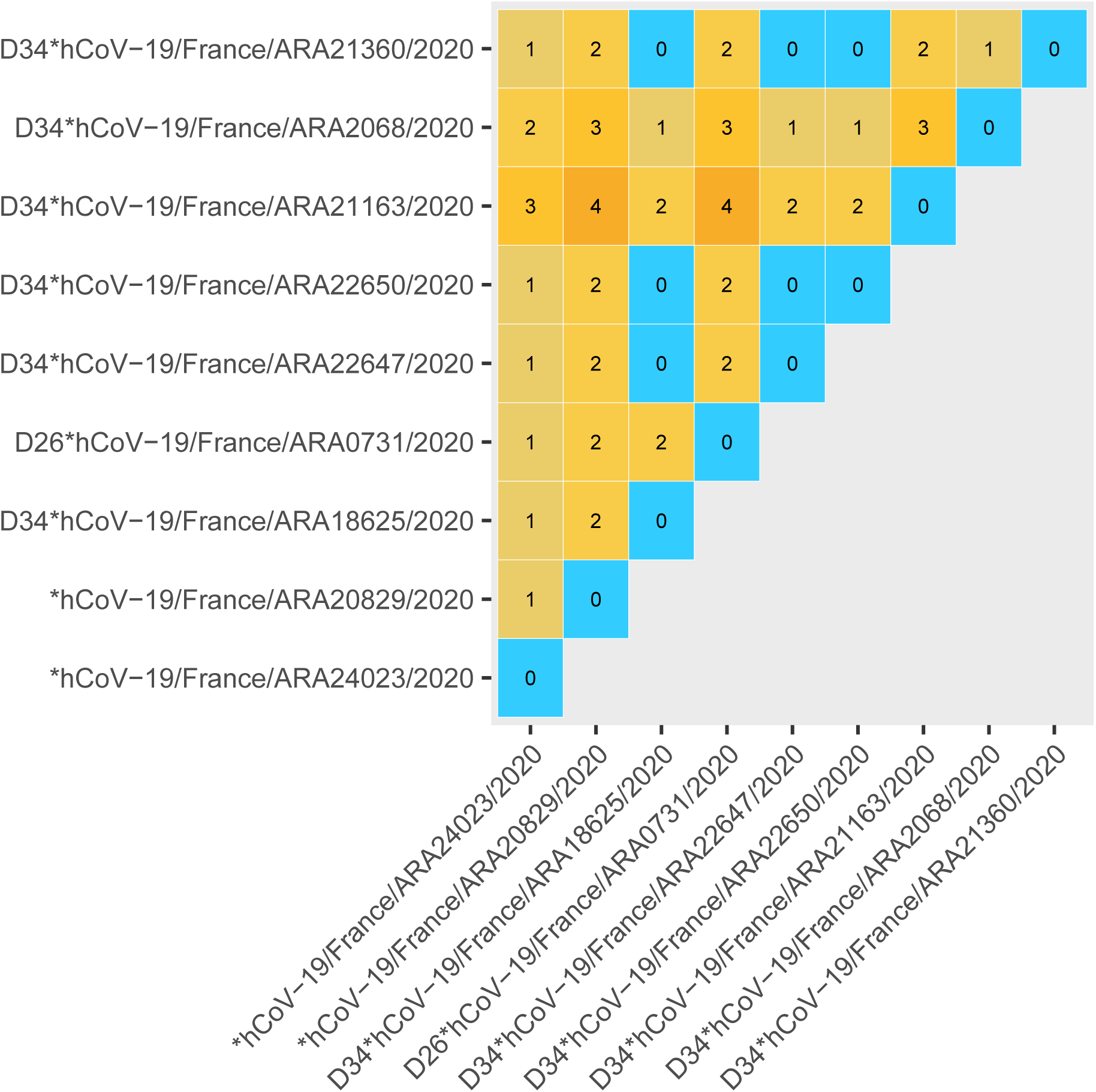
Single nucleotide polymorphism heat map of SARS-CoV-2 ORF6 WT and deletion strains. Mismatch count between consensus sequences generated by each method compared 2 by 2 for each sample. Blue tiles correspond to perfect identity and orange tiles correspond to mismatches (number of mismatches is indicated inside the tile). Matrices were generated with an R script using Decipher (alignment), ape (distance matrices) and ggplot2 (charts) libraries. Of note, undetermined bases and deletions were notconsidered in the calculation of mismatches.

**SUPPLEMENTARY TABLE 1.**
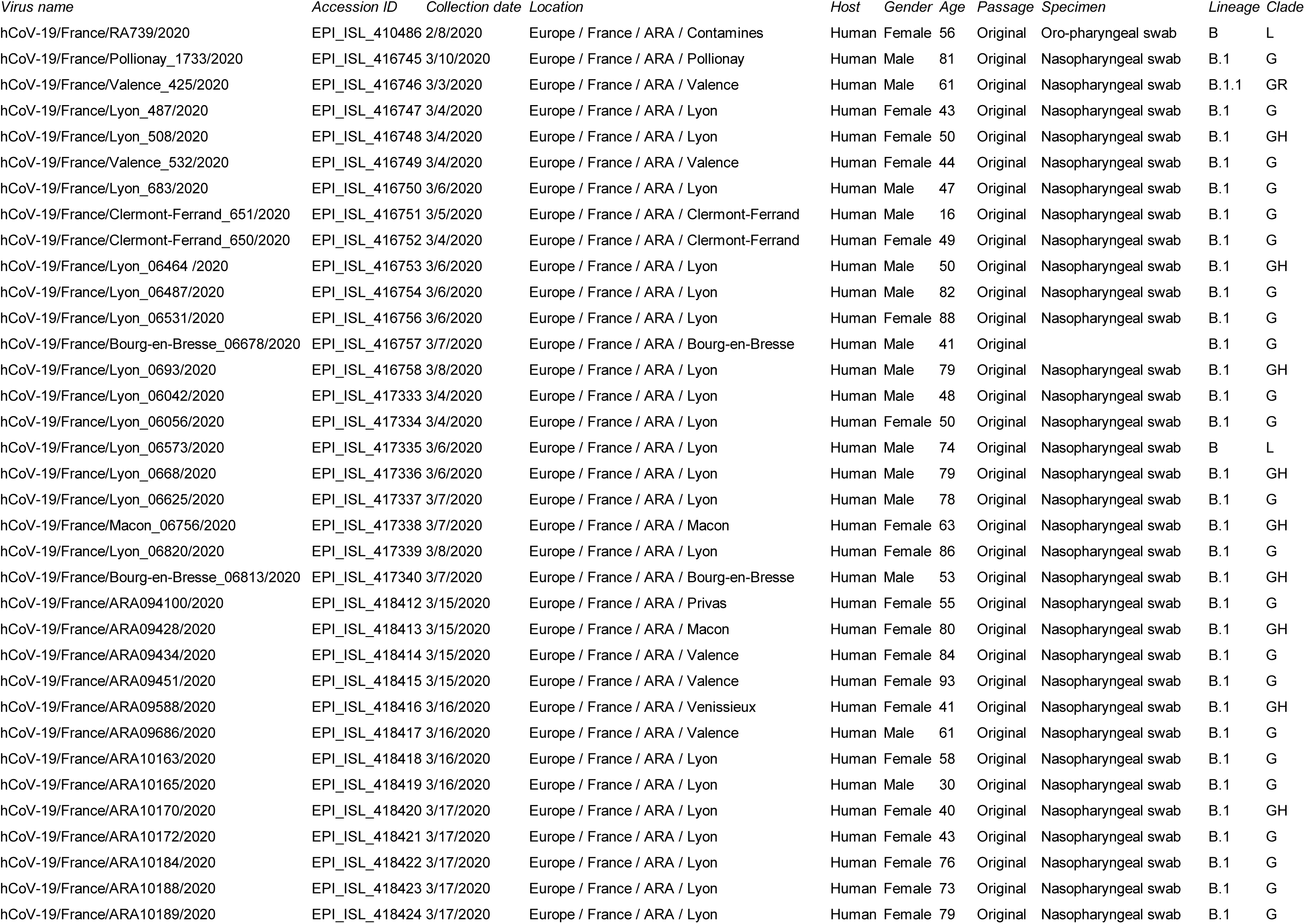
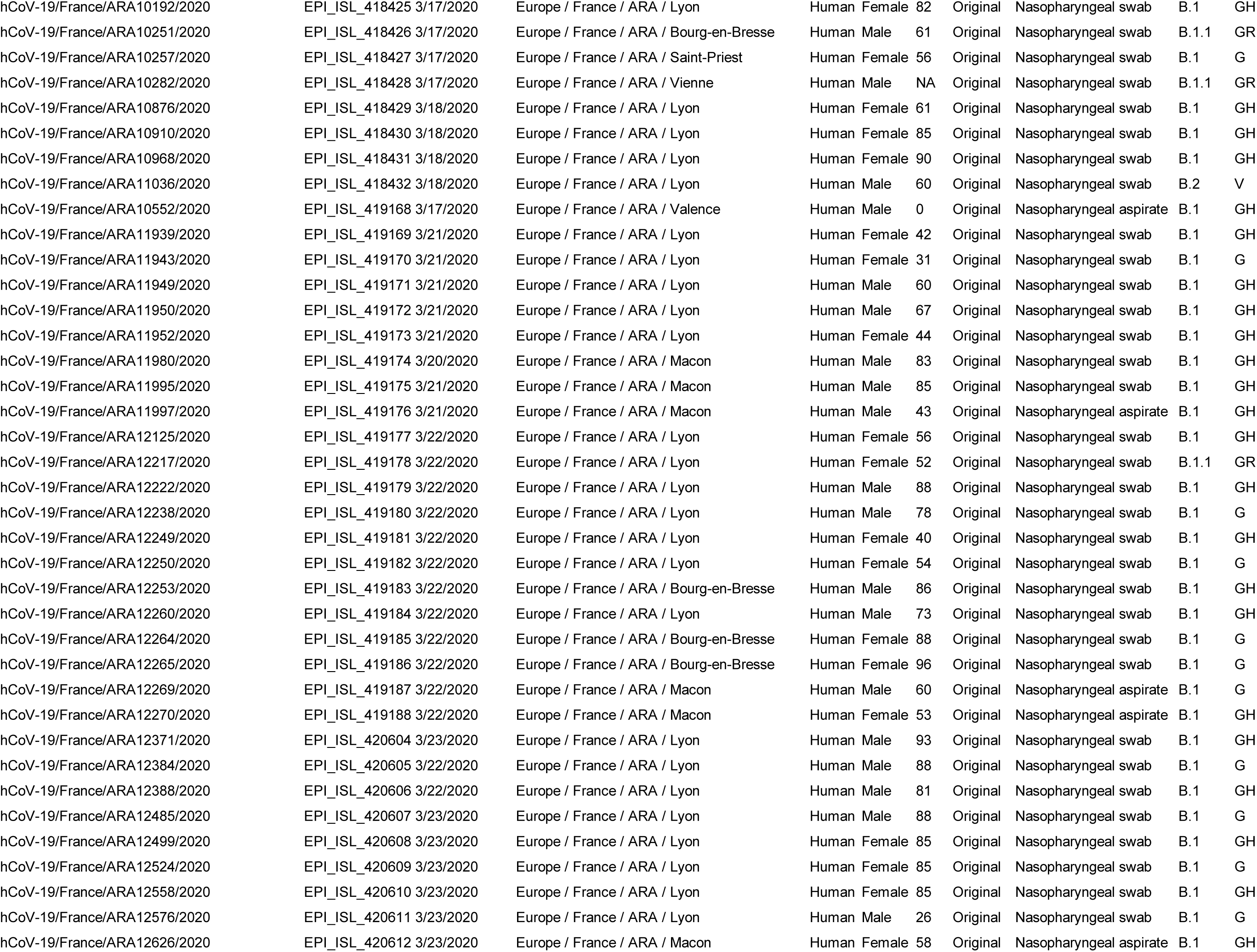
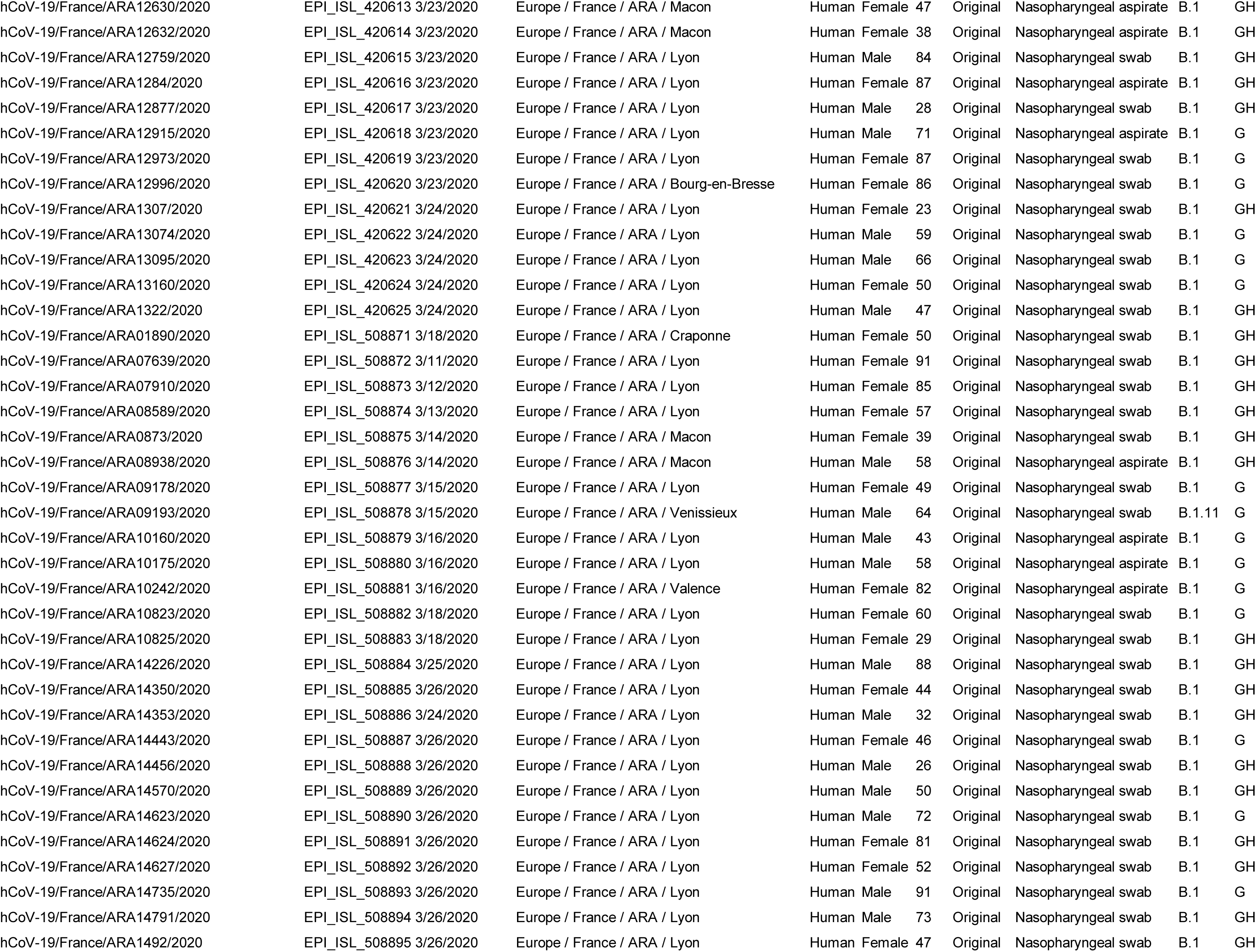
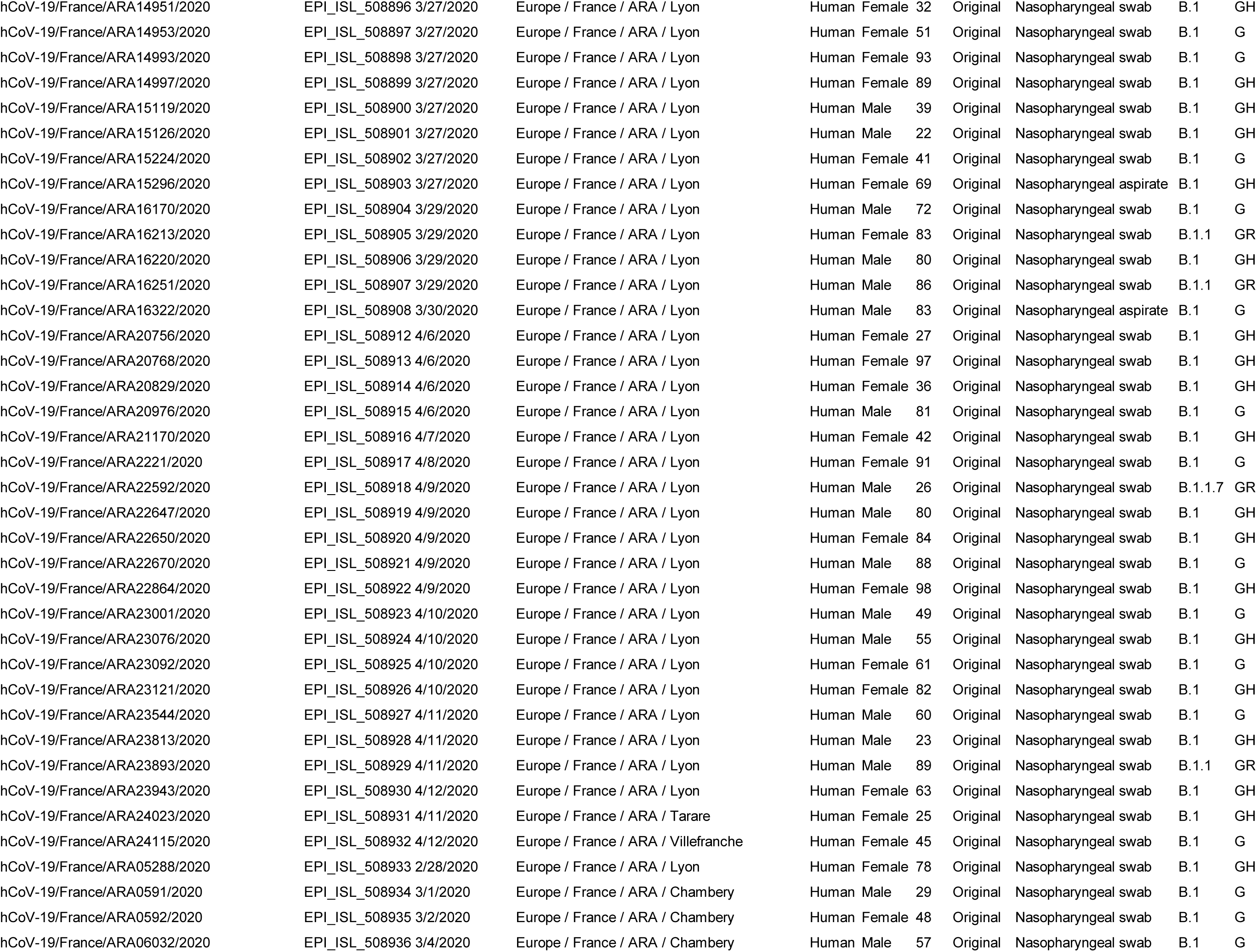
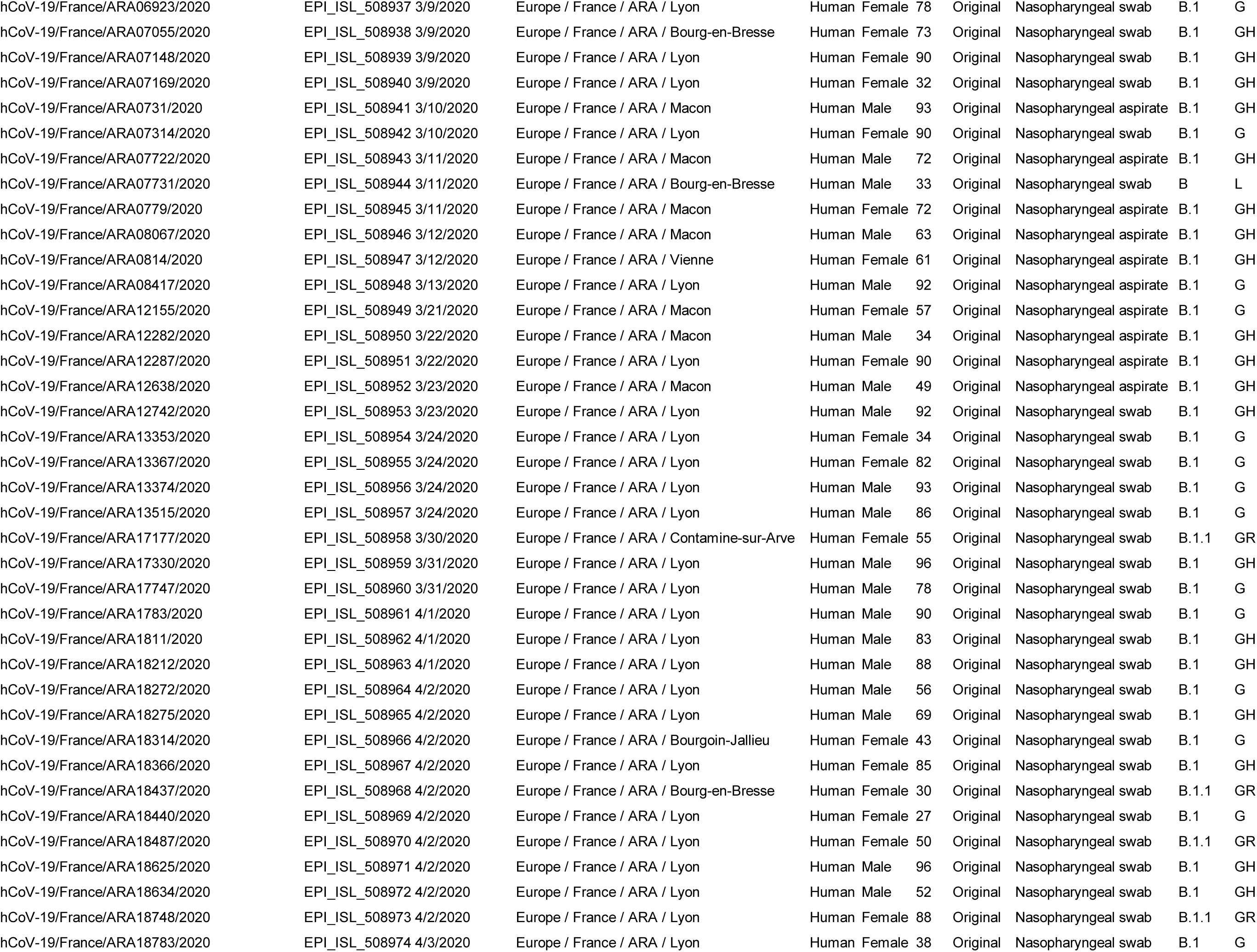
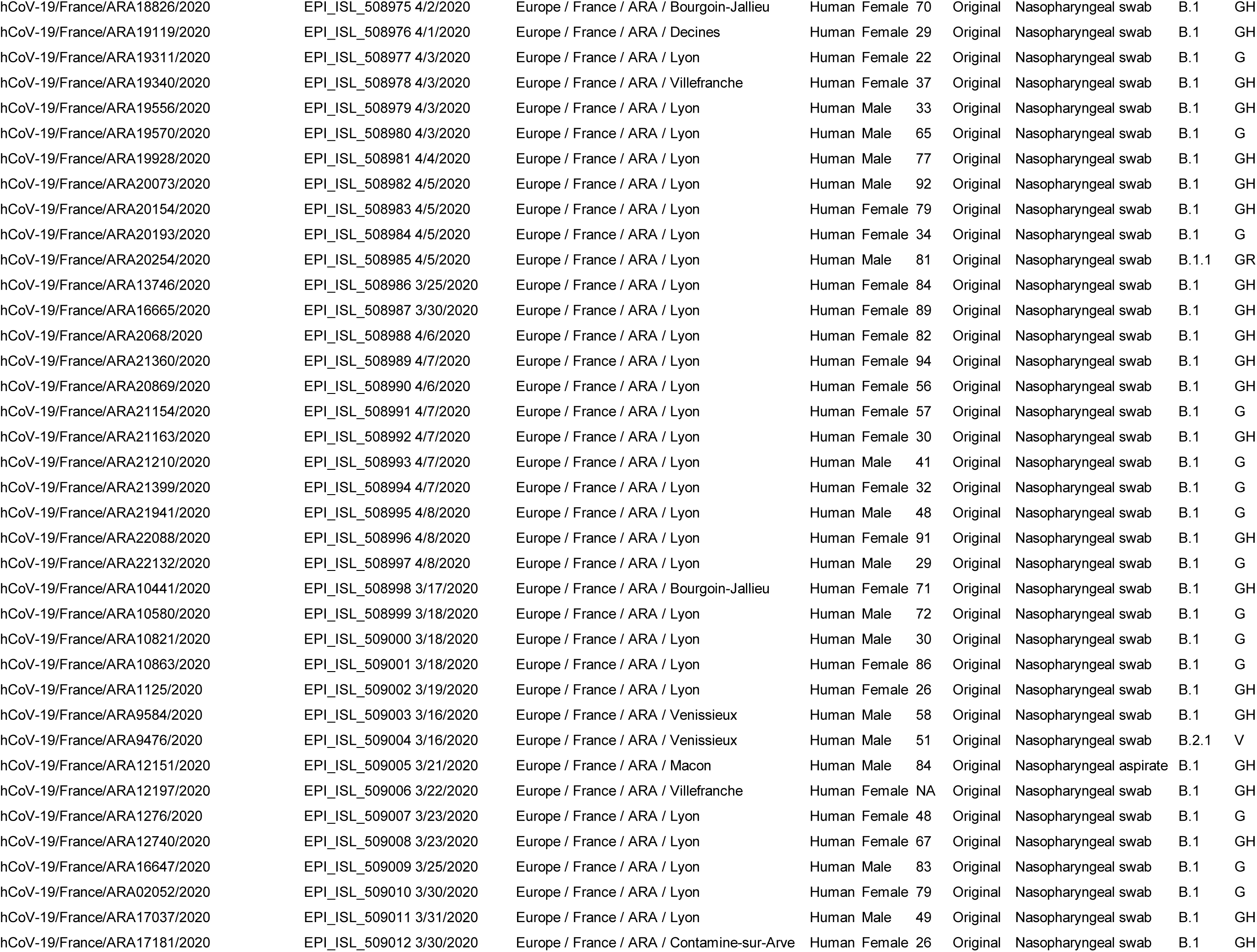
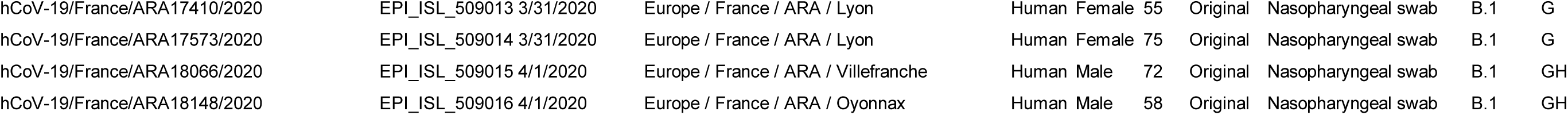
SARS-CoV-2 genomes sequenced in the Auvergne-Rhônes-Alpes region from Feb 12 to Apr 12, 2020

